# The acute effects of cannabidiol on emotional processing and anxiety: a neurocognitive imaging study

**DOI:** 10.1101/2020.12.11.421776

**Authors:** Michael AP Bloomfield, Yumeya Yamamori, Chandni Hindocha, Augustus PM Jones, Jocelyn LL Yim, Hannah R Walker, Ben Statton, Matthew B Wall, Rachel H Lees, Oliver D Howes, H Valerie Curran, Jonathan P Roiser, Tom P Freeman

## Abstract

**Background:** There is growing interest in the therapeutic potential of cannabidiol (CBD) across a range of psychiatric disorders. CBD has been found to reduce anxiety during experimentally-induced stress in anxious individuals and healthy controls. However, the mechanisms underlying the putative anxiolytic effects of CBD are unknown. We therefore sought to investigate the behavioural and neural effects of a single dose of CBD vs. placebo on a range of emotion-related measures to test cognitive-mechanistic models of its effects on anxiety.

**Methods:** We conducted a randomised, double-blind, placebo-controlled, crossover, acute oral challenge of 600 mg of CBD in 24 healthy participants on emotional processing, with neuroimaging (viewing emotional faces during fMRI) and cognitive (emotional appraisal) measures as well as subjective response to experimentally-induced anxiety.

**Results:** CBD did not produce effects on BOLD responses to emotional faces, cognitive measures of emotional processing, or modulate experimentally-induced anxiety, relative to placebo.

**Conclusions:** Given the rising popularity of CBD for its putative medical benefits, further research is warranted to investigate the clinical potential of CBD for the treatment of anxiety disorders.

---

Anxiety disorders constitute a leading global cause of morbidity (World Health Organization, 2017) and are associated with considerable economic burden (Revicki et al., 2012). In the search for novel treatments, cannabidiol (CBD) is a cannabinoid that has been proposed to possess anxiolytic effects which may be clinically promising across a range of anxiety disorders (Blessing, Steenkamp, Manzanares, & Marmar, 2015). Research into CBD’s potential anxiolytic properties is much needed to stimulate progress in novel pharmacological approaches to treatment, given only moderate response rates of around 50-60% (for example in cognitive behavioural therapies (Loerinc et al., 2015) and serotonin reuptake-inhibitors (Baldwin, Woods, Lawson, & Taylor, 2011)), and major cutbacks in psychiatric drug development (Abbott, 2011; Miller, 2010).

CBD elicits anxiolytic effects comparable to those of classical anxiolytic drugs (e.g. diazepam) in animal anxiety models, including the marble-burying test (Casarotto, Gomes, Resstel, & Guimaraes, 2010), the Vogel conflict test (Moreira, Aguiar, & Guimaraes, 2006) and the elevated plus maze (Guimaraes, Chiaretti, Graeff, & Zuardi, 1990). Despite this, and in contrast with standard drugs, CBD’s effects are not dependent on serotonergic or GABAergic signalling as CBD’s anxiolytic effects were neither blocked by a 5HT_1A_ receptor antagonist (Casarotto et al., 2010) nor a benzodiazepine antagonist (Moreira et al., 2006). However, CBD’s effects on fear learning were blocked by the endocannabinoid receptor type 1 (CB1R) antagonist, SR141716A (Bitencourt, Pamplona, & Takahashi, 2008), which suggests that anxiolysis is attributable to the endocannabinoid system. This endocannabinoid pathway has been proposed as a novel pharmacological target for anxiety (Hill & Gorzalka, 2009; Lutz, Marsicano, Maldonado, & Hillard, 2015; Ruehle, Rey, Remmers, & Lutz, 2012) and therefore cannabinoid research may provide new means to treat these disorders.

In humans, the anxiolytic effects of CBD have been supported through tasks involving anxiogenic public speech paradigms in healthy individuals (Linares et al., 2019; Zuardi, Cosme, Graeff, & Guimaraes, 1993), socially anxiety (Bergamaschi et al., 2011), in clinical high risk of psychosis (Appiah-Kusi et al., 2020) and in Parkinson’s disease (de Faria et al., 2020). Further, CBD also reduced drug cue-induced anxiety in heroin-abstinent individuals (Hurd et al., 2019). The effects of CBD on emotional processing, i.e. the cognitive processing of emotion-related information, have also been investigated. CBD appears to facilitate extinction learning of fear memories in humans (Das et al., 2013), improve performance in the recognition of ambiguous emotional facial expressions (Hindocha et al., 2015) and shift attention away from emotional stimuli to more neutral stimuli (Arndt & de Wit, 2017). Functional magnetic resonance imaging (fMRI) studies have shown attenuated amygdala and anterior cingulate responses (as well as attenuated connectivity between these regions) to fearful faces following CBD administration (Fusar-Poli et al., 2010; Fusar-Poli et al., 2009) and this effect was correlated with reduced subjective anxiety (Bhattacharyya et al., 2010). One finding complicates this general trend as CBD produced a non-significant increase in anxiety ratings in a paranoid sample during virtual reality (Hundal et al., 2018). Yet, the majority of these results support the anxiolytic hypothesis of CBD, which may be mediated by the neurocognitive effects of CBD on emotional processing.

Despite recent research into the anxiolytic role of CBD, little attention has been given to the neurocognitive mechanisms underlying CBD’s effects on anxiety – that is, whether CBD’s neural effects and emotional processing modulation constitute mechanisms of this anxiolytic effect. Thus, we sought to assess CBD-induced anxiolysis from a neurocognitive perspective to investigate candidate mechanisms. Firstly, we measured CBD’s effect on the neural correlates of emotional processing in an fMRI emotion viewing task, with a region-of-interest in the amygdala. Secondly, we measured CBD’s behavioural effects on emotional appraisal using a task adapted from a previous study which was sensitive to the effects of CBD in reward processing (Hindocha et al., 2018). Thirdly, we measured CBD’s effects on anxiety responses to experimentally-induced stress. Lastly, we explored the relationship between these measures to assess the link between CBD’s effects on neurocognitive mechanisms of emotional processing and its effects on subjective anxiety. Specifically, we hypothesised that, compared to placebo, CBD would: i) attenuate automatic responses of the amygdala to negative emotions; ii) lower valence and arousal scores in appraising facial expressions of both positive and negative valence; and lastly that CBD would iii) attenuate experimentally-induced anxiety in the stress task.

## Experimental procedures

We used a double-blind, placebo-controlled, crossover design to assess the effects of acute CBD challenge in healthy participants. The order of CBD and placebo administration was randomised and balanced for sex. The research procedure was approved by the UCL Research Ethics Committee (reference: 3325/002) and conducted in accordance with the Helsinki Declaration of 1975.

### Participants

Twenty-four healthy participants (12M, 12F) were recruited. Inclusion criteria were: i) healthy volunteer; ii) English-speaking; iii) age 18-70 years; iv) right-handed. Primary exclusion criteria included lifetime CBD use, psychiatric history and fMRI contraindications (see the supplementary methods for full exclusion criteria). All participants gave written informed consent. A sensitivity analysis conducted with G*Power 3.1.9.2. (Faul, Erdfelder, Lang, & Buchner, 2007) indicated that our sample size provided 84% power to detect a significant difference (*p* < .05, two-tailed) between CBD and placebo with an estimated effect size of *d* = 0.5, based on previous findings of acute CBD on subjective ratings of drug-related stimuli (*d* = 0.5; Hindocha et al., 2018).

### Drugs

Participants were administered 600 mg of oral CBD (pure synthetic (−)-CBD, STI Pharmaceuticals, Essex, United Kingdom) or matched placebo (lactose powder) in 12 identical and opaque capsules in each testing session. The dose of 600 mg was selected given previous evidence of CBD’s anxiolytic effect at this dose (Bergamaschi et al., 2011), and a waiting period of 2.5 h before testing was chosen based on of evidence for peak plasma CBD around this time (Haney et al., 2016). We collected blood samples via venepuncture 4 h post-drug administration (after MRI) in EDTA vacutainers for immediate centrifugation. Plasma samples were stored at −80 °C and analysed using gas chromatography and mass spectrometry with a lower limit of quantification of 0.5ng/ml.

### Behavioural measures

#### Subjective/physiological measures

We recorded state measures of mood (‘happy’, ‘anxious’) at 5 time-points across each session, using 11-point VAS anchored at 0 (‘not at all’) to 10 (‘extremely’). Heart rate (HR) and systolic and diastolic blood pressure (SBP, DBP) were also measured. Measurements were made 10 min before drug administration (baseline) and subsequently at 0.5 h, 2 h, 4 h, and 6 h post-drug administration. We used the Beck Anxiety Inventory (Beck, Epstein, Brown, & Steer, 1988) to measure trait anxiety symptoms over the previous week.

#### Face rating task

This task measured subjective emotion appraisal (based on the pleasantness rating task in Hindocha et al., 2018). Stimuli were male and female adult open-mouth happy/angry/neutral expressions from the NimStim set of facial expressions (Tottenham et al., 2009). The actors were changed across sessions. Participants viewed the faces in randomised order and made valence and arousal judgments on each face. The valence judgment was described through the question: “How positive or negative does the image look to you?” and rated along a 7-point VAS from “-3: very negative” to “+3: very positive”. The arousal judgment was described through the question: “How aroused does the image make you feel?” and rated along a 7-point VAS from “0: not at all aroused” to “+6: extremely aroused”. Faces remained on the screen until both rating judgments were made. Participants were instructed to respond as quickly and accurately as possible, and that arousal referred to emotional reaction rather than sexual arousal.

#### Mental arithmetic task

A mental arithmetic task (Constantinou et al., 2010) was implemented to measure emotional responses to stress. The task was comprised of two parts: a no-stress (control) condition and a stress condition, in fixed order (no-stress, stress) to avoid carry-over stress effects to the no-stress condition. *Part 1: No-stress condition.* Participants were given a paper handout consisting of a series of arithmetic additions and subtractions. They were asked to simply circle the character ‘3’ every time it appeared for 3 min. Participants were told that speed and accuracy were not important. *Part 2: Stress condition (stress task).* Participants were asked to verbally and serially subtract 13s from 6022 (session 1) or 6021 (session 2) as quickly and as accurately as they could for 3 min. Participants were prompted on incorrect responses and reminded of the importance of accuracy and speed after the 1st and 2nd minute to promote stress regardless of performance. Verbal responses were recorded using an external microphone held before the participant, although these recordings were not analysed. HR, SBP, DBP, and VAS scores of ‘anxious’, ‘stressed’, ‘calm’, and ‘relaxed’ were recorded four times throughout the task: immediately after the control task instructions, immediately after completing the control task, immediately after the stress task instructions, and immediately after completing the stress task (pre-control, post-control, pre-stress, and post-stress, respectively).

### Neuroimaging measures

#### fMRI paradigm

The task was based on a previous incidental emotional processing task (O’Nions, Dolan, & Roiser, 2011). The participants were shown a series of faces and asked to categorise them on the basis of sex (male/female) by key-press. There were eight cycles of three 16 s blocks (happy, fearful, and neutral) in fixed order. Each block was composed of eight faces representing one emotion, with each face presented for 1500 ms followed by a central fixation cross of 500 ms. There was a 2 s interval between each block, and 16 s interval of rest following each cycle of three blocks. The stimuli were 42 faces also obtained from the NimStim set of facial expressions (Tottenham et al., 2009). Faces were presented in pseudo-random order such that there were equal numbers of male and female faces. As task performance was irrelevant to the main measure of automatic emotional processing, the behavioural data are not discussed further (see supplementary information for MRI acquisition and pre-processing procedures).

### Procedure

Participants attended two sessions with at least 7 days between sessions to minimise carry-over drug effects (Haney et al., 2016; Hindocha et al., 2018). We asked participants to fast from midnight (12 am) on the night before each session, based on a previous study (Haney et al., 2016). Water was permitted anytime thereafter and a caffeinated drink for participants routinely consuming caffeine was also permitted to avoid caffeine withdrawal. We also asked participants to avoid alcohol for 24 h before sessions and to avoid smoking on the morning of sessions. Pre-drug checks consisted of breath tests for alcohol and tobacco (via carbon monoxide) use, drug urine screens and pregnancy tests for female participants. Drug administration (0 h) was followed by the waiting period, after which fMRI scanning occurred (2 h 30 min post-drug). Participants were then provided a standard meal followed consisting of a sandwich, snack and drink (4h). Finally, the behavioural tasks were administered (5 h 30 min post-drug). Other measures were recorded as part of the procedure however these are reported elsewhere (Bloomfield et al., 2020; Lawn et al., 2020). The procedure for the present study lasted approximately 7 h (Table 1).

**Table 1.** Timeline for each testing session.

| **Time from drug administration** | **Measure** |
| --- | --- |
| - 10 min | Physiological and subjective measures (baseline) |
| + 0 h | Drug administration - cannabidiol or placebo |
| + 30 min | Physiological and subjective measures (T1) |
| + 2 h | Physiological and subjective measures (T2) |
| + 2 h 30 min | fMRI scanning (emotional processing task) |
| + 4 h | Blood samples, physiological and subjective measures (T3) |
| + 5 h 30 min | Behavioural tasks (face rating, mental arithmetic) |
| + 6 h | Physiological and subjective measures (T4) |

*Note.* We recorded physiological and subjective measures including subjective anxiety at 5 timepoints over the course of a single testing session (baseline, T1-T4). All procedures were time-locked to the time of drug administration. Other neuroimaging and behavioural measures were employed as part of the study, however these are reported elsewhere.

### Statistical analyses

Behavioural data were performed with frequentist analysis via Statistical Package for the Social Sciences 25 (SPSS Inc., Released 2020.) and Bayesian analysis via JASP (JASP Team, 2020). Frequentist and Bayesian repeated-measures analyses of variance (ANOVAs) were performed, comparing the effect of drug (CBD, placebo) and other task factors (e.g. emotion, time) on each task’s dependent variables. Bayes factors were computed from the Bayesian repeated-measures ANOVAs, which measure the relative predictive performance of the null and alternative hypotheses, which in turn allowed us to evaluate the strength of evidence in favour of the null hypothesis (which is assumed to be true, but not tested, in frequentist analyses). Post-hoc tests were conducted through a priori orthogonal contrasts relevant to the task variables: contrasting happy vs. neutral and angry vs. neutral in the face rating task, and as Helmert contrasts in the mental arithmetic task and the periodic subjective/physiological measures. Outliers, defined as data points which were more than three times the interquartile range from the nearest quartile, were winsorised to the nearest non-outlier value. We re-tested all statistical models with drug administration order as a between-subjects factor to account for practice/familiarity effects. A significance threshold of *ɑ* = .05 (2-tailed) was adopted for the frequentist analyses. For the Bayesian analyses, we followed standard guidelines for interpreting the Bayes factors in accordance with Jeffreys (1998).

The neuroimaging (MRI) statistical model was specified by creating three regressors of happy, fearful and neutral affect. Realignment motion parameters were included as covariates-of-no-interest. A high-pass filter of 128 s was applied. Neural responses to positive and negative emotion were modelled by contrasts of happy *vs.* neutral faces, and fearful *vs.* neutral faces, respectively. Task and drug effects were analysed across the whole brain at a cluster-level threshold of *ɑ* = .05 (FWE-corrected), and through an anatomically-defined amygdala region-of-interest (ROI) at a within-ROI voxel-level threshold of *ɑ* = .05 (FWE-corrected). A voxel-based ROI analysis was preferred over single-parameter extraction, as multiple amygdala sub-regions were used to produce the ROI with potentially distinct sensitivities for positive and negative emotion. The ROI was built using the SPM Anatomy Toolbox (Eickhoff et al., 2005), by conglomerating masks of the basolateral, centromedial and superficial sub-regions (Amunts et al., 2005) into a single mask of the bilateral amygdala. The SPM Anatomy Toolbox was also used to localise activation maps.

## Results

The inclusion of order as a between-subjects factor also did not affect any drug-related findings, so this was not included in the final models.

### Sample characteristics

Four participants were excluded due to: aversion to MRI (n = 2), gastro-intestinal discomfort following lunch (n = 1) and positive test for tricyclic antidepressant use (n = 1). The final sample consisted of 24 participants (12 male, 12 female), and had a mean age of 23.6 years (SD = 4.12), BMI of 22.3 kg/m^2^ (SD = 3.48), and low BAI score of 2.6 (SD = 3.23) which is in the range of ‘normal to no anxiety’ (Julian, 2011). All participants had a Fagerström Test for Nicotine Dependence score (Heatherton, Kozlowski, Frecker, & Fagerstrom, 1991) of 0, whilst the mean score on the Alcohol Use Disorders Identification Test (Saunders, Aasland, Babor, de la Fuente, & Grant, 1993) was 2 (SD = 2.1). The mean interval between sessions was 9.5 days (SD = 4.25).

### Blood plasma CBD concentration

Analysis of plasma CBD levels via Wilcoxon signed-rank test revealed higher CBD plasma concentrations in CBD sessions (median = 6.01 ng/ml, IQR = 4.24) compared to placebo (median = 0 ng/ml, IQR = 0) sessions (*z* = 4.10, *p* < .001, *r* = .88).

### Neuroimaging results

#### Effect of task

For the happy *vs.* neutral faces contrast, we found significantly increased BOLD response in the right calcarine gyrus (*p*_FWE-corrected_ < .001; Figure 1A, Table 2A). For the fearful *vs.* neutral faces contrast, we found significantly increased response in the left lingual gyrus (*p*_FWE-corrected_ < .001; Figure 1B, Table 2B). We did not find any significant reductions in BOLD response across the whole brain for either contrast. Within the amygdala ROI, we found a non-significant effect of the fear *vs.* neutral faces contrast in the right basolateral amygdala (*p*_FWE-corrected_ = .057; Figure 1C, Table 2B*).

**Figure 1.**
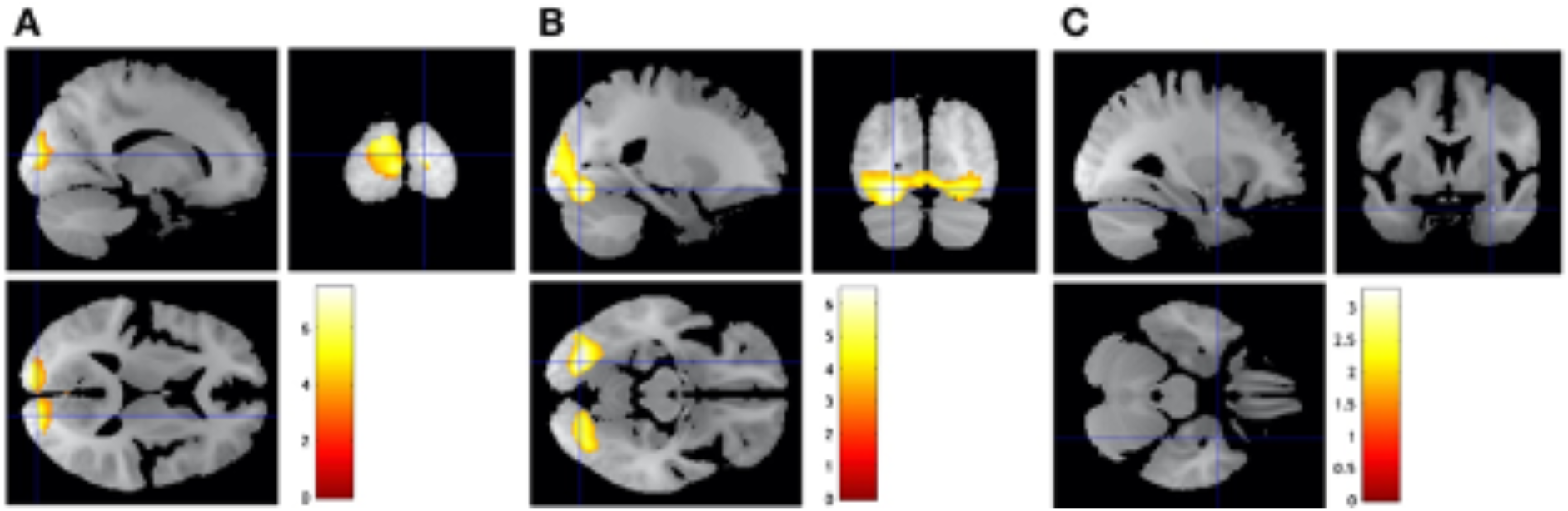
A) Happy faces, relative to neutral faces, increased BOLD responses in the right calcarine gyrus. B) Fearful faces, relative to neutral faces, increased BOLD responses in the left lingual gyrus. C) There was a non-significant effect within the a priori amygdala region of interest. Statistical maps are overlaid on the sample mean structural image. A voxel-based threshold of ɑ = .001 (uncorrected) was used to form the beta clusters, after which a cluster-level threshold of ɑ = .05 (FWE-corrected) was applied. Coloured bars indicate t-values.

**Table 2.** Regions with significant increases in BOLD response during the fMRI emotional processing task across the whole brain and *a non-significant finding within the amygdala region-of-interest (ROI)

| Contrast | Region | MNI coordinates (mm) |  |  | Cluster size (mm<sup>3</sup>) | Z-score |
| --- | --- | --- | --- | --- | --- | --- |
|  |  | x | y | z |  |  |
| a. happy vs neutral | Right Calcarine Gyrus | 16 | -92 | 11 | 13776 | 6.02 |
| b. fearful vs neutral | Left Lingual Gyrus | -22 | -78 | -13 | 24468 | 5.45 |
|  | *Right Amygdala | 30 | 2 | -25 | 12 | 3.10 |

A voxel-based threshold of $\alpha = .001$ (uncorrected) was used to form the beta clusters, after which a cluster-level threshold of $\alpha = .05$ (FWE-corrected) was applied. \*This finding was under the FWE-corrected threshold of $\alpha = .05$ at $p_{\text{FWE-corrected}} = .057$ .

**Figure 2.**
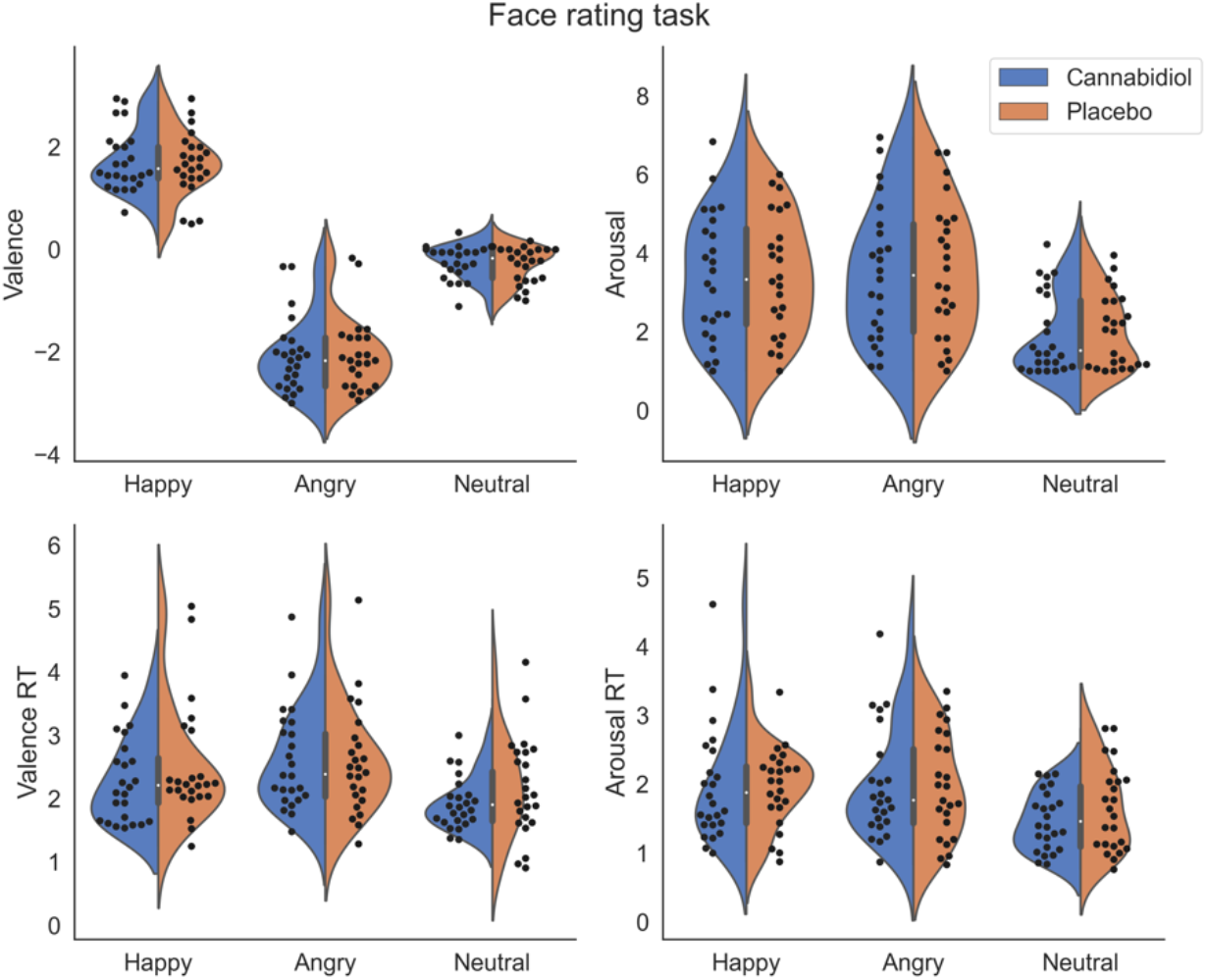
Subjective valence and arousal scores from the face rating task and their reaction times. There was no evidence for differences across drug (cannabidiol vs placebo).

**Figure 3.**
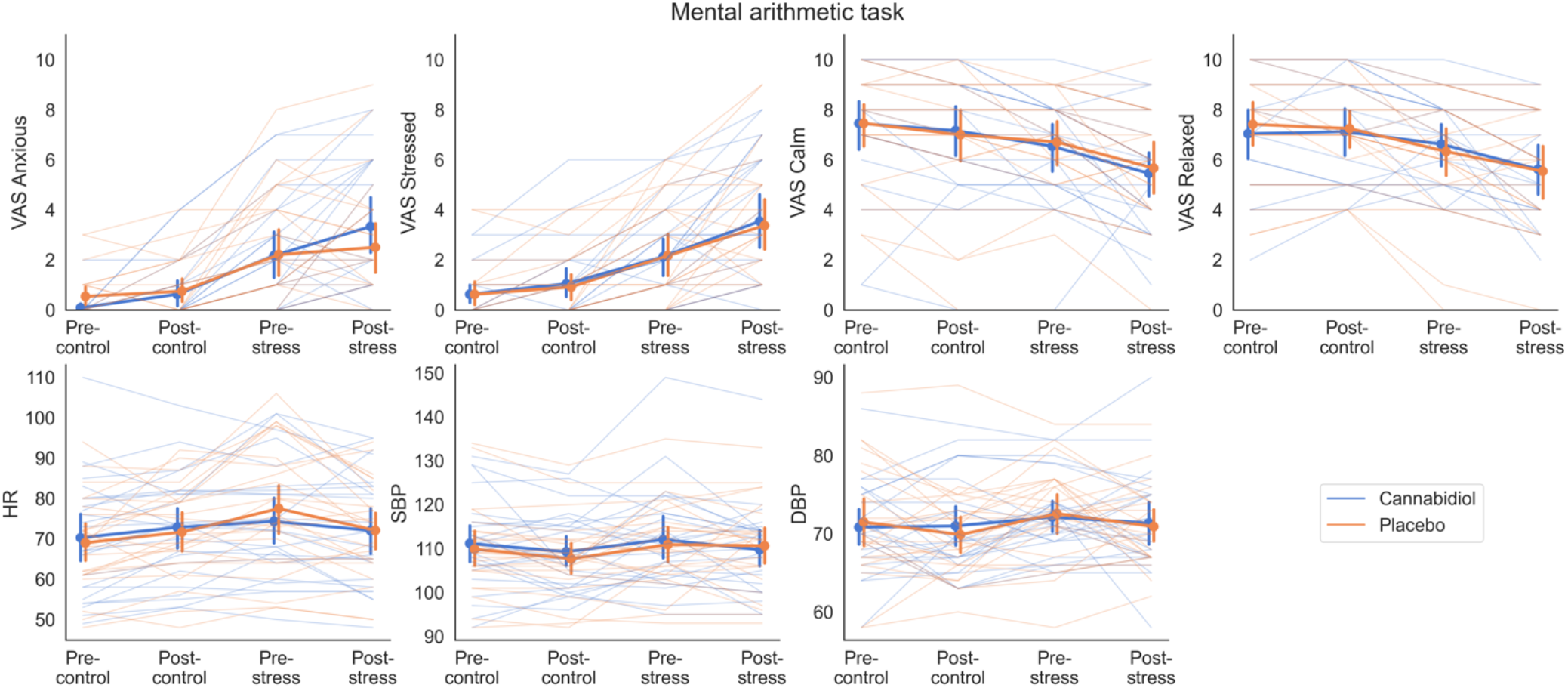
Subjective and physiological responses during the mental arithmetic task. Frequentist analyses suggested an interaction effect of drug and time on subjective anxiety (p = .048) where the increase in pre-stress to post-stress was greater for cannabidiol vs placebo, however this was contradicted by Bayesian analysis (BF_10_ = 0.387). No other effects were evident. Error bars represent 95% confidence intervals.

**Figure 4.**
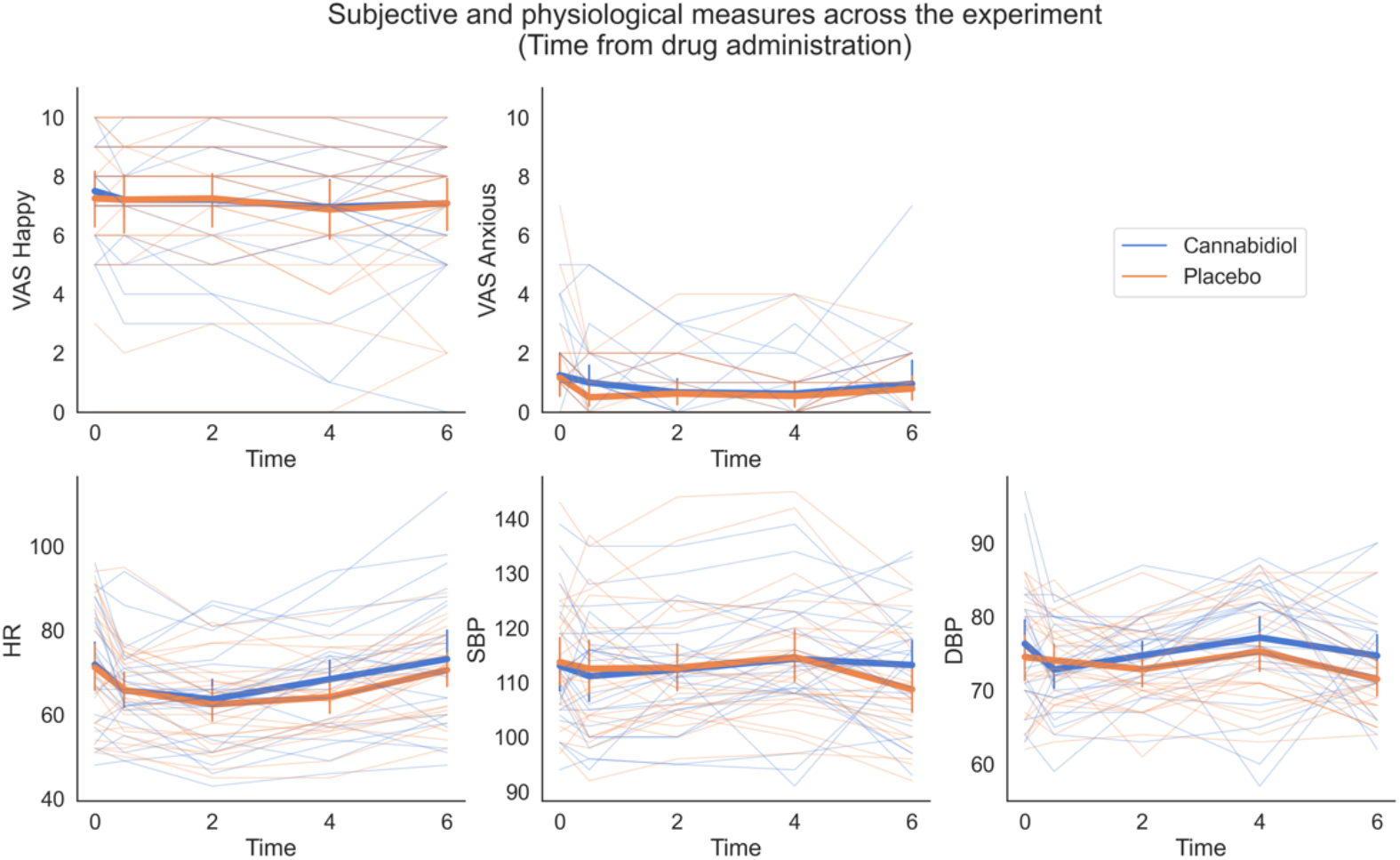
Subjective and physiological measures across the duration of the experiment (time from drug administration). There was no evidence for differences across drug (cannabidiol vs placebo). Error bars represent 95% confidence intervals.

#### Effect of drug

We did not find significant drug effects for either the happy *vs.* neutral faces, or the fearful *vs.* neutral faces contrasts across the whole-brain and within the ROI.

### Behavioural results

Drug-relevant behavioural effects (main effects of drug and drug-related interactions) are reported below (see supplementary results for full statistical analyses).

### Face rating task

We did not find significant drug-related effects for valence (drug: *F*_1,23_ = 0.54, *p* = .468, BF_10_ = 0.180; drug x emotion: *F*_1.39,31.93_ = 0.16, *p* = .771, BF_10_ = 0.118) and arousal (drug: *F*_1,23_ = 0.29, *p* = .597, BF_10_ = 0.183; drug x emotion: *F*_1.26,29.01_ < 0.01, *p* = .975, BF_10_ = 0.115) judgments, and similarly for valence (drug: *F*_1,23_ = 0.98, *p* = .333, BF_10_ = 0.504; drug x emotion: *F*_2,46_ = 2.67, *p* = .080, BF_10_ = 0.220) and arousal (drug: *F*_1,23_ = 0.26, *p* = .617, BF_10_ = 0.228; drug x emotion: *F*_1.43,32.81_ = 1.13, *p* = .318, BF_10_ = 0.181) RTs.

### Mental arithmetic task

#### VAS ‘anxious’ and ‘stressed’

We did not find a main effect of drug for anxiety (*F*_1, 23_ = 0.07, *p* = .799, BF_10_ = 0.155). Frequentist analysis showed a significant interaction effect of drug and time (*F*_2.14, 49.29_ = 3.17, *p* = .048, *η*^*2*^*p* = .12), although this was not supported by Bayesian analysis (BF_10_ = 0.387) which rather suggested anecdotal evidence towards the null hypothesis, and suggests that the frequentist finding represents a Type I error. We did not find significant drug-related effects for stress (drug: *F*_1,23_ = 0.06, *p* = .802, BF_10_ = 0.157; drug x time: *F*_2.09,48.11_ = 0.11, *p* = .904, BF_10_ = 0.062).

#### VAS ‘calm’ and ‘relaxed’

We did not find significant drug-related effects for calm (drug: *F*_1,23_ = 0.04, *p* = .846, BF_10_ = 0.164; drug x time: *F*_2.04,46.91_ = 0.36, *p* = .705, BF_10_ = 0.070) and relaxed (drug: *F*_1,23_ = 0.02, *p* = .899, BF_10_ = 0.154; drug x time: *F*_1.91,43,92_ = 1.14, *p* = .327, BF_10_ = 0.115) ratings.

#### Physiological measures

We did not find significant drug-related effects for heart rate (drug: *F*_1,23_ = 0.02, *p* = .901, BF_10_ = 0.156; drug x time: *F*_3,69_ = 2.26, *p* = .089, BF_10_ = 0.212), systolic blood pressure (drug: *F*_1,23_ = 0.48, *p* = .496, BF_10_ = 0.247; drug x time: *F*_3,69_ = 0.79, *p* = .503, BF_10_ = 0.108) or diastolic blood pressure (drug: *F*_1,23_ = 0.02, *p* = .887, BF_10_ = 0.158; drug x time: *F*_3,69_ = 1.18, *p* = .322, BF_10_ = 0.107).

### Subjective and physiological measures

We did not find significant drug-related effects for subjective anxiety (drug: *F*_1,23_ = 0.69, *p* = .416, BF_10_ = 0.318; drug x time: *F*_2.74,63.08_ = 0.54, *p* = .639, BF_10_ = 0.054) or happiness (drug: *F*_1,23_ = 0.08, *p* = .785, BF_10_ = 0.163; drug x time: *F*_2.21,50.82_ = 0.24, *p* = .805, BF_10_ = 0.037) across the experiment. Similarly, we did not find significant drug-related effects for heart rate (drug: *F*_1,23_ = 2.45, *p* = .128, BF_10_ = 0.610; drug x time: *F*_2.46,56.58_ = 1.09, *p* = .354, BF_10_ = 0.094), systolic blood pressure (drug: *F*_1,23_ = 0.14, *p* = .717, BF_10_ = 0.151; drug x time: *F*_3.05,70.15_ = 1.98, *p* = .124, BF_10_ = 0.328) or diastolic blood pressure (drug: *F*_1,23_ = 2.77, *p* = .110, BF_10_ = 1.640; drug x time: *F*_4,92_ = 1.51, *p* = .204, BF_10_ = 0.187).

## Discussion

The results suggest that CBD did not have any effects on our measures relating to anxiety. Whilst we confirmed that CBD was absorbed into blood plasma, contrary to our hypothesis, CBD did not modulate neural or behavioral correlates of emotional processing via incidental emotion viewing, or emotion appraisal, and had no effect on subjective and cardiovascular responses to experimentally-induced stress. These null findings follow those of other studies reporting minimal behavioural effects of CBD (Arndt & de Wit, 2017; Babalonis et al., 2017; Haney et al., 2016).

We may have observed our negative findings as a result of a lack of sensitivity of our measures to CBD’s effects. Evidence against this possibility is that we selected our measures on the basis of their previous sensitivity to drug-related effects (Hindocha et al., 2018; Selvaraj et al., 2018) and that each of our measures elicited significant task effects (e.g. differentiation of valence and arousal responses to positive and negative emotional stimuli in the face rating task, increased subjective and cardiovascular stress responses in the mental arithmetic task, see the supplementary materials for full statistical results). This pattern of results mirror those of Ardnt and de Wit (2017) who also found no effects of CBD on a range of emotion-related measures despite significant task effects.

Critically, our null findings for the effect of CBD on experimentally-induced stress are in contrast to previous findings reporting that CBD attenuated task-induced anxiety (Bergamaschi et al., 2011; Zuardi et al., 1993; Zuardi et al., 2017), and these discrepant findings may be due to the potentially dose-dependent nature of CBD’s effects. Recent work suggests that CBD may have an inverted-U shape dose-response curve (Campos & Guimaraes, 2008; Freeman et al., 2020; Hsiao, Yi, Li, & Chang, 2012; Linares et al., 2019; Zuardi et al., 2017) with best efficacy for human anxiety at 300 mg (Linares et al., 2019; Zuardi et al., 1993; Zuardi et al., 2017) compared to our dose of 600 mg. However, the finding that CBD was able to reduce drug-cue-induced craving and anxiety at single doses of 400 mg and 800 mg (Hurd et al., 2019), suggests that our dose of 600 mg still falls within the effective range, and so our negative finding remains important as evidence against the anxiolytic hypothesis of CBD.

Lastly, three previous studies have found that chronic administration of CBD was anxiolytic: a 21-day treatment of 600 mg of CBD was effective at reducing psychosis-related anxiety in patients at high-risk of psychosis, compared to placebo (Bhattacharyya et al., 2018) and a four-week treatment of oil with 300 mg of CBD significantly reduced anxiety in a socially anxious sample, compared to placebo (Masataka, 2019). Additionally, four-week treatment with 800mg CBD reduced anxiety in people with a cannabis use disorder compared to placebo (Freeman et al., 2020). Therefore, repeated dosing of CBD may necessary to produce anxiolytic effects.

### Strengths and limitations

There are several strengths of the present study. Our experimental design allowed for a simultaneous test of CBD’s effects across neurocognitive and subjective levels of emotional processing. We used previously validated measures (Constantinou et al., 2010; O’Nions et al., 2011), especially with respect to the cognitive tasks which were sensitive to the effects of CBD in a reward processing context (Hindocha et al., 2018). Further, the measures tested for both positive and negative-emotional processing, whereas previous studies have only focused on the latter, especially in neuroimaging studies.

The limitations of the study are that the oral route of CBD administration is slow and associated with variable bioavailability. For example, Haney et al. (2016) reported that oral administration of 800 mg resulted in a wide spread of peak concentrations of CBD in plasma from 1.6 to 271.9 ng/ml, and the times of peaks of CBD plasma varied from 120 to 360 minutes. We also employed a relatively long fasting time compared to other studies (overnight vs 2 hours), which may have been detrimental as co-administering CBD with food can increase oral bioavailability (Taylor, Gidal, Blakey, Tayo, & Morrison, 2018). However, our plasma measures showed that there was significant absorption of CBD. Additionally, the stimuli employed across tasks were inconsistent; the fMRI task used fearful faces to represent negative emotion, whilst the face rating task used angry faces.

Finally, one of our measures was novel in the context of CBD research, which was the mental arithmetic task, whereas previous studies have employed public speaking (Appiah-Kusi et al., 2020; Bergamaschi et al., 2011; Zuardi et al., 1993; Zuardi et al., 2017) and virtual reality (Hundal et al., 2018) paradigms. Importantly, unlike the public speaking task, an advantage of this mental arithmetic task is its suitability for use in a repeated-measures design. Yet, since we did not find CBD-related effects in any of our measures, it is impossible to determine whether the mental arithmetic task was less sensitive than previous tasks, or other factors contributed to CBD’s null effect.

## Conclusions

The present study found no effect of a single dose of 600 mg CBD on a range of emotional processing measures in a healthy sample, despite multiple previous reports of anxiolytic effects of CBD and effects of CBD on emotional processing. These findings warrant further investigation in light of increasing popularity of CBD and its potential use for treating anxiety disorders.

## Acknowledgements

We are extremely grateful to all the radiographers at the Robert Steiner MRI Unit and Professor Glyn Lewis. We would also like to thank all our participants for taking part in this study. We are grateful to Ting-Yun Chang for her help with preparing this manuscript.

## Ethical standards

The authors assert that all procedures contributing to this work comply with the ethical standards of the relevant national and institutional committees on human experimentation and with the Helsinki Declaration of 1975, as revised in 2008.

## Contributors

Authors MABP, TPF, & CH designed the study and wrote the protocol. Author YY managed the literature searches. Authors YY, APMJ, JLLY, HRW, RHL & BS coordinated the study and conducted assessments. Authors YY & MBW conducted the analyses. Authors MABP and YY wrote the first draft of the manuscript. All authors contributed to and have approved the final manuscript.

