## Supplementary materials for "The acute effects of cannabidiol on emotional processing and anxiety: a neurocognitive imaging study"

Full inclusion and exclusion criteria

Inclusion criteria were: i) healthy volunteer; ii) English-speaking; iii) age 18-70 years; iv) right-handed. Exclusion criteria were: i) current use of psychotropic drugs; ii) current/past use of cannabis or cannabidiol; iii) no more than 5 occurrences of recreational drug use other than cannabis; iv) current or history of mood, psychotic, anxiety or substance abuse disorder assessed with an adapted version of the structured clinical interview for DSM-IV (First, Gibbon, Spitzer, Williams, & Benjamin, 1997) (SCID-IV); v) current nicotine dependence defined by a score greater than 3 on the Fagerström Test for Nicotine Dependence (Heatherton, Kozlowski, Frecker, & Fagerstrom, 1991) (FTND); vi) hazardous/harmful alcohol use defined by a score greater than 7 on the Alcohol Use Disorders Identification Test (Saunders & Babor, 1993) (AUDIT); vii) pregnancy; viii) lack of capacity; ix) needle phobia; x) colour blindness, xi) allergies to/unwillingness to take cannabidiol, microcrystalline cellulose, gelatin or lactose; and xii) contraindications to fMRI.

MRI acquisition and pre-processing procedures

***MRI acquisition.*** Scanning was performed on a Siemens 3T Prisma MRI scanner. A total of 181 whole-brain volumes of 44 axial slices were collected with a T2*-weighted echo-planar imaging (EPI) sequence (TR = 3000 ms, TE = 30 ms, slice thickness = 3 mm, 2 x 2 mm in-plane resolution, phase encoding direction = anterior → posterior, field of view = 250 mm2, matrix size = 64 x 64, flip angle = 90°). A whole-brain structural volume of 176 axial slices was acquired afer the functional scans with a T1-weighted magnetisation-prepared rapid gradient-echo (MPRAGE) sequence (TI = 900 ms, TR = 2300 ms, TE = 2.28 ms, slice thickness = 1 mm, 1 mm2 in-plane resolution, phase encoding direction = anterior → posterior, field of view = 256 mm2, matrix size = 256 x 256, flip angle = 9°).

***Pre-processing.*** The fMRI data were pre-processed using MATLAB (The Mathworks Inc., 2017) and Statistical Parametric Mapping (Friston, Ashburner, Kiebel, Nichols, & Penny, 2007) (SPM12). The first four volumes of each functional scan were discarded to allow for T1 equilibration. The scans were then realigned to the new first image using a least-squares approach and a 6-parameter (rigid-body) affine transformation. The dimensions of the functional image voxels were used as translation thresholds (i.e. 2 mm for x and y translations, 3 mm for z translations), and 2° was used as the rotation threshold. No images exceeded these motion thresholds. The scans were normalised to the Montreal Neurological Institute- (MNI) 152 template, using a 12-parameter affine transformation, and smoothed using a 8 x 8 x 8 mm full-width half-maximum Gaussian kernel.

Non-drug-related statistical results

A priori outliers

All behavioural models were re-tested after the exclusion of a) the participant who had breakfast, and b) the two participants who were outside healthy BMI ranges, to determine whether these a priori outliers affected any drug-related effects (main effects of drug or drug interaction effects). As this did not affect any results, the full sample was retained for all tests. Drug administration order as a between-subjects factor also did not affect the results, so are not discussed further.

Since the drug-related results are reported in the main article, here we only report the effects which did not include a factor of drug (i.e. effects only relating to task factors).

Face rating task

*Valence.* With respect to valence judgments, there was a significant main effect of emotion (*F_1.16,26.73_* = 263.17, *p* < .001, *η^2^p* = .92, BF_10_ = 2.170x10^64^). Post-hoc tests revealed a significant difference across all conditions (*p*_Holm_ < .001), where happy faces were rated as more positive than neutral faces, and neutral faces were rated as more positive than angry faces. With respect to RTs, there was a main effect of emotion (*F_2,46_* = 12.55, *p* < .001, *η^2^p* = .35, BF_10_ = 549). Post-hoc tests showed greater RTs for happy compared to neutral faces (*p*_Holm_ = .005), and similarly greater for angry compared to neutral faces (*p*_Holm_ < .001). This effect was likely due to the VAS design of responses, where neutral responses did not require as much movement of a visual pointer and therefore time for each response.

*Arousal.* With respect to arousal judgments, there was a main effect of emotion (*F_2, 46_* = 15.12, *p* < .001, *η^2^p* = .40, BF_10_ = 2.317x10^8^). Post-hoc tests revealed significant differences in ratings between happy and neutral, and angry and neutral conditions (*p*_Holm_ < .001), where happy and angry faces were rated as more arousing compared to neutral faces. With respect to RTs, there was a main effect of emotion (*F_2,46_* = 15.77, *p* < .01, *η^2^p* = .41, BF_10_ = 948). Post-hoc tests showed greater RTs for happy compared to neutral faces (*p*_Holm_ = .005), and similarly greater for angry compared to neutral faces (*p*_Holm_ < .001). which were again likely due to the VAS design of responses.

Mental arithmetic task

*VAS ‘anxious’ and ‘stressed’.* With respect to anxiety, there was a significant main effect of time (*F_1.67,38.32_* = 24.36, *p* < .001, *η^2^p* = .51, BF_10_ = 5.061x10^14^), where repeated contrasts showed that anxiety significantly increased from post-control to pre-stress (*p* < .001) and from pre-stress to post-stress (*p* = .043). With respect to stress, there was a significant main effect of time (*F_1.49,34.29_* = 28.82, *p* < .001, *η^2^p* = .56, BF_10_ = 1.804x10^16^) where repeated contrasts showed that stress significantly increased from post-control to pre-stress, and from pre-stress to post-stress (*p* < .001).

*VAS ‘calm’ and ‘relaxed’.* With respect to calmness, there was a main effect of time (*F_1.72,39.59_* = 16.23, *p* < .001, *η^2^p* = .41, BF_10_ = 5.954x10^7^), where repeated contrasts showed that calmness decreased from pre-stress to post-stress (*p* < .001). A similar pattern of results was observed for relaxedness: there was a main effect of time (*F_1.61,37.07_* = 16.40, *p* < .001, *η^2^p* = .42, BF_10_ = 7.756x10^7^), where repeated showed that relaxedness significantly decreased from post-control to pre-stress scores (*p* = .011), and pre-stress to post-stress scores (*p* = .001).

*Physiological measures.* With respect to SBP, there was a significant main effect of time (*F_3,69_* = 2.88, *p* = .042, *η^2^p* = .11, BF_10_ = 0.662), where repeated contrasts showed that SBP significant increased from post-control to pre-stress (*p* = .005). There was no effect of time for DBP (*F_3,69_* = 1.90, *p* = .138, BF_10_ = 0.357). With respect to HR, there was a significant main effect of time (*F_3, 69_* = 8.08, *p* < .001, *η^2^p* = .26, BF_10_ = 186). Inspection of the repeated contrasts showed that HR significantly changed across adjacent measurements: HR increased from pre-control to post-control (*p* = .046), increased from post-control to pre-stress (*p* = .006) and decreased from pre-stress to post-stress (*p* = .004).

Subjective and physiological measures

*VAS ‘anxious’ and ‘happy.* A main effect of time was observed (*F_2.91,66.94_* = 3.36, *p* = .025, *η^2^p* = .13, BF_10_ = 1.040), where anxiety significantly decreased from baseline to 1 h post-drug administration (*p* = .018). There was no effect of time for ‘happy’ scores (*F_2.80,64.33_* = 2.28, *p* = .092, BF_10_ = 0.163).

*Physiological measures.* There was no effect of time for SBP (*F_2.72,62.52_* = 2.15, *p* = 108, BF_10_ = 0.391). A significant main effect of time was observed for DBP (*F_4,92_* = 3.99, *p* = .005, *η^2^p* = .15, BF_10_ = 4.814), where DBP changed between each time-point except for between 1 and 2 h post-drug administration (*p* < .05). A main effect of time on HR was also observed (*F*_4,92_ = 14.98, *p* < .001, *η^2^p* = .39, BF_10_ = 6.389x10^8^), where HR changed between each time point except for between 1 and 2 h post-drug administration (*p* < .05).
